# A Novel Role for CSA in the Regulation of Nuclear Envelope Integrity: Uncovering a Non-Canonical Function

**DOI:** 10.1101/2023.12.14.571633

**Authors:** Denny Yang, Austin Lai, Amelie Davies, Anne FJ Janssen, Delphine Larrieu

**Affiliations:** Pharmacology Department, University of Cambridge, Tennis Court Road CB2 1PD Cambridge, United Kingdom; Cambridge Institute for Medical Research, The Keith Peters Building, Hills Road CB2 0XY Cambridge, United Kingdom; Duke University Medical Center, Durham, Trent Drive 27710 NC United States

## Abstract

Cockayne syndrome (CS) is an autosomal recessive premature ageing condition mainly characterized by microcephaly, growth failure, and neurodegeneration. It is caused by mutations in *ERCC6* or *ERCC8* genes which encode for Cockayne Syndrome B (CSB) and Cockayne Syndrome A (CSA) proteins, respectively. CSA and CSB have well-characterised roles in transcription-coupled nucleotide excision repair (TC-NER), responsible for the removal of bulky DNA lesions, including those caused by UV irradiation. Here, we report that CSA knockout cells and CSA patient cells (CS-A) carrying a loss-of-function mutation in the *ERCC8* gene exhibit defects in nuclear envelope (NE) integrity. NE dysfunction is a characteristic phenotype of cells from progeroid disorders caused by mutation in NE proteins, such as Hutchinson-Gilford Progeria Syndrome (HGPS). However, it has never been reported in Cockayne Syndrome. We observed that CS-A cells displayed reduced levels of LAP2-emerin-MAN1 (LEM)-domain 2 (LEMD2) at the NE resulting in decreased formation of LEMD2-lamin A/C complexes. In addition, loss of CSA function caused increased actin stress fibers that contributed to enhanced mechanical stress to the NE. Altogether, these led to NE blebbing and ruptures in interphase, causing activation of the innate/immune cGAS/STING signaling pathway. Disrupting the linker of the nucleoskeleton and cytoskeleton (LINC) complex that is responsible for anchoring the cytoskeleton to the NE, rescued the NE phenotypes and reduced the activation of cGAS/STING pathway. This work has revealed a previously uncharacterized role for CSA in regulating NE integrity and shed light on mechanisms that may further explain some of the clinical phenotypes observed in CS patients such as neuroinflammation. This is to our knowledge, the first study showing NE dysfunction in a progeroid syndrome caused by mutations in a DNA damage repair protein, reinforcing the connection between NE deregulation and ageing.

## Introduction

Progeroid disorders are a group of incurable rare diseases that resemble many aspects of physiological aging (1,2). Clinical manifestations include premature and accelerated ageing, developmental delay, hair loss, vision and hearing loss, progressive neurodegeneration, cardiovascular diseases, osteoporosis, and early death (1). Progeroid disorders can be characterized by two distinct subtypes at the molecular level. The conditions caused by mutations in genes involved in the DNA damage response and repair, including Werner Syndrome, Ataxia Telangiectasia, Bloom Syndrome, Cockayne Syndrome and Xeroderma Pigmentosum, and the conditions caused by mutations in nuclear envelope (NE) genes, such as Hutchinson Gilford Progeria Syndrome (HGPS) or Nestor Guillermo Progeria Syndrome (NGPS) (3).

Cockayne syndrome (CS), is an autosomal recessive condition caused by mutations in either *ERCC8* (20% of cases) or *ERCC6* (80% of cases) genes, encoding for CSA and CSB proteins, respectively (4–6). CS occurs at a rate of 1 in 300,000-500,000 live births in the USA and Europe, and the patients have a life expectancy ranging from 5 to 16 years (7,8). CS is mainly characterized by growth failure and neurological abnormalities (9). Other clinical manifestations include cataracts, microcephaly, and cutaneous photosensitivity (10). The main described function for CSA and CSB proteins is in transcription-coupled nucleotide excision repair (TC-NER). TC-NER is a DNA damage repair mechanism that removes bulky DNA adducts such as the ones induced by UV, mainly including 6-4 pyrimidine-pyrimidine (6-4 PP) photoproducts and cyclobutene-pyrimidine dimers (CPD) (11,12). The loss-of-function mutations in *ERCC8* and *ERCC6* genes occurring in Cockayne Syndrome patients, prevent the removal of these bulky DNA lesions by TC-NER (12,13). This results in the progressive accumulation of DNA damage in CS patient cells, causing cellular photosensitivity and explaining some of the clinical symptoms in CS patients (5,14). However, other clinical manifestations of CS such as neurodegeneration (15) cannot be explained by defects in TC-NER, since other TC-NER-associated syndromes including UV sensitive syndrome (UVSS) do not display neurodegeneration (3,16). This suggests that there are additional mechanisms underlying tissue dysfunction in CS patients. New functions for CS proteins in mitochondrial autophagy and oxidative stress response (17,18), as well as in transcriptional regulation have been suggested to contribute to these phenotypes (15).

The NE plays a pivotal function in regulating cellular homeostasis, by maintaining the structural architecture of the nucleus and by controlling chromatin organization, nucleocytoplasmic transport, and propagation of mechanical cues from the cytoplasm to the nucleus (19–21). The NE is composed of two lipid bilayers, the inner and outer nuclear membranes (INM and ONM). The ONM is continuous with the endoplasmic reticulum (ER) and merges with the INM at the nuclear pore complexes. The nuclear lamina is a protein meshwork composed of intermediate filaments of A-type lamin (lamin A and C) and B-type lamin (lamin B1 and B2) proteins (22). lamins interact with the chromatin and with other NE proteins including LAP2-emerin-MAN1 (LEM)-domain proteins (23). Extracellular mechanical signals are sensed and propagated to the nucleus through the linker of the nucleoskeleton and cytoskeleton (LINC) complex (24), composed of SUN-domain proteins (SUN1 and SUN2) and KASH-domain proteins (Nesprin-1-4) (25). The LINC complex spans over the INM and ONM connecting cytoskeletal components to the nucleus. Loss of NE integrity can be triggered by various stresses.

NE integrity can be challenged by deleterious mutations in genes encoding for NE proteins (e.g. *LMNA* or *LMNB1)* (26) or by mechanical stress (e.g. cancer cells migrating through tiny blood vessels during metastasis or tissues subjected to contractions such as skeletal muscles or heart) (27–29). These stresses enhance the probability for the cells to experience NE ruptures during interphase. These ruptures are typically preceded by the formation of gaps in the lamina that generate a weak point at the NE. This leads to the formation of nuclear blebs in which the chromatin can protrude through the lamina gaps (30,31). Nuclear blebs can then rupture if the mechanical pressure is not resolved. This results in exchange of content between the nucleus and the cytoplasm which can result in DNA damage and activation of the innate immune cGAS/STING pathway, known to be associated with inflammation (32). The NE repair process is mediated by Barrier-to-Autointegration Factor 1 (BAF) where LEM domain proteins, A-type lamins and endosomal sorting complexes required for transport (ESCRT)-III protein complex are recruited to the rupture site to facilitate the resealing of the nuclear membrane and restore NE integrity (31,33–35).

Here, we found that CSA knockout or loss-of-function was associated with multiple NE abnormalities, causing ruptures and activation of the cGAS/STING innate immune pathway. More specifically, we revealed two mechanisms that drive NE defects in CSA patient cells: (1) the decreased formation of LEMD2-lamin A/C complexes at the NE and (2) the increased actin stress fibers that generate mechanical tension on the NE. This work sheds light on a new role for CSA in NE regulation and suggests a new mechanism that can contribute to loss of homeostasis in CS-A cells.

## Material and Methods

### Cell lines and cell culture

The CSA patient-derived fibroblast cells (CS3BE, termed CS-A in this study), CSB patient-derived fibroblast cells (CS1AN, termed CS-B in this work), and their respective isogenic cell lines (complemented with HA-CSA - WT(HA-CSA) or HA-GFP-CSB - WT(HA-GFP-CSB), were a kind gift from Dr. Sebastian Iben. Cells were grown in complete medium consisting of Dulbecco’s Modified Essential Medium with 10% (v/v) fetal bovine serum (FBS) (Gibco) and 1% (v/v) penicillin/streptomycin (10,000 U/mL) (Thermo-Fisher). The complemented “WT” cells were grown in complete media with 50 ug/mL of Geneticin (Life Technologies) as a selection media to maintain HA-CSA or HA-GFP-CSB plasmid expression.

CSA and CSB knockout HAP1 cell lines were purchased from Horizon Discovery and cultured in Iscove’s Modified Essential Medium media, containing 10% FBS and 1% (v/v) penicillin/streptomycin (10,000 U/mL).

### Drug treatments

For drug treatments, cells were seeded onto 12-well plate at 50% confluency. Drug concentration and treatment time used throughout this study are outlined in Table 1. At the start of each drug treatment, culture media was replaced with fresh media containing the working concentration of the indicated drugs **(Table 1)**.

**Table 1:** Details of chemical used for drug treatment experiments.

| Drug | Working Concentration | Incubation time/hour | Manufacturer | Catalog Number |
| --- | --- | --- | --- | --- |
| Cytochalasin D | 0.25 µg/mL | 18 hours | Cambridge Bioscience | 11330 |
| Jasplakinolide | 25 nM | 6 hours | Abcam | ab141409 |

### Plasmid transfection and constructs

Cells were first seeded onto tissue culture plate at 70% confluency. Next day, transfection of plasmid constructs was performed using Trans-IT 2020 reagent (Mirus Bio, #MIR 5400) following the manufacturer’s protocols. 24 hours after transfection, experiments were performed. pEGFP-C1 plasmid construct was purchased from Clontech. Flag-CSA construct subcloned in pcDNA3.1 vector was a kind gift Dr. Sebastian Iben. LEMD2-GFP was a kind gift from Dr. Kyle Roux (33) and was subcloned into the pEGFP-C1 plasmid.

### siRNA knockdown

The SUN1 siRNA oligonucleotide was purchased from Sigma-Aldrich with the following sequence: siSUN1: 5’-CCAUCCUGAGUAUACCUGUCUGUAUDTDT-3’, siLEMD2: 5’-UUGCGGUAGACAU CCCGGGDTDT-3’. MISSION® siRNA Universal Negative Control #1 (Sigma-Aldrich, #SIC001) was used as the control siRNA. Cells were first seeded onto tissue culture plate at 50-60% confluency. The next day, the transfection of siRNA was performed using Lipofectamine RNAiMAX (Thermo Fisher, #13778075) following the manufacturer’s instructions. 30 pmol of siRNA were transfected into cells in 12 well plates for 48 hours.

### Western blot

Protein extraction from monolayered culture cells was performed by scraping cells in SDS lysis buffer (4% SDS, 20% glycerol, 120 mM Tris-HCl, pH = 6.8), boiling for 5 minutes at 95°C, and passing through a 25-gauge needle 10 times. For samples requiring pre-extraction of soluble proteins, cells were first incubated with cold cytoskeletal buffer (CSK) (100 mM NaCl, 300 mM sucrose, 1 mM EGTA, 1 mM MgCl_2_, 1 mM DTT, 10 mM PIPES/KOH, 6.8 pH) for 5 minutes on ice. Protein concentration was measured using the nanodrop spectrophotometer (Thermo-Fisher) at 280 nm. 30 μg of each protein sample was then heat-denatured 5 min at 95°C after adding Protein Sample Loading Buffer (Li-cor, #928-40004) with 100 mM dithiothreitol (DTT). Denatured protein samples and PageRuler^TM^ Prestained Protein ladder (Thermo-Fisher, #26616) and PageRuler^TM^ Plus Prestained Protein ladder (Thermo-Fisher, #26619) was size separated using precast NuPAGE^TM^ 4-12%, Bis-Tris, 1.0-1.5 mm, Mini Protein Gels (Thermo-Fisher, #NP0322BOX) in 1X NuPAGE™ MES SDS Running Buffer (Thermo-Fisher, #NP0002) at 180V. Size-separated proteins were then transferred onto a nitrocellulose membrane in 1X NuPAGE^TM^ Transfer Buffer (Thermo-Fisher, #NP0006) at 250 mA. SDS-PAGE and protein transfer were performed using the Mini Gel Tank (Thermo-Fisher, #A25977) and Mini Blot Module (Thermo-Fisher, #B1000) system, respectively. The nitrocellulose membrane was blocked with 5% (w/v) non-fat milk dissolved in Tris-buffered saline (TBS, 50 mM Tris-HCl, 150 mM NaCl) with 0.1% (v/v) Tween-20 (Sigma-Aldrich) (0.1% TBS-T) for 1 hour at room temperature with gentle agitation. Primary antibody incubation was performed for either 1 hour at room temperature or overnight at 4°C. Secondary antibody incubation was performed for 1 hour at room temperature. Details of primary and secondary antibodies used are outlined in **Table 2**. Protein detections were carried out using the Odyssey CLx Imager (LI-COR) system. For densitometric analysis, band intensities of the protein of interest were normalized to the band intensity of housekeeping protein and followed by normalizing all tested samples to the WT or control samples.

**Table 2:** Details of primary and secondary antibodies used for immunofluorescence and western blot experiments.

| Primary Antibodies |  |  |  |  |
| --- | --- | --- | --- | --- |
| Antibody | Western Blot Dilution | Immunofluorescence Dilution | Supplier | Catalogue Number |
| Anti-LEMD2 | 1:200 | 1:200 | Sigma | HPA017340 |
| Anti-SUN1 | 1:1000 | 1:100 | Abcam | ab124770 |
| Anti-alpha-tubulin | 1:500 | 1:500 | Sigma | T9026 |
| Anti-cGAS | 1:250 | 1:250 | Cell Signalling | D1D3G |
| Anti-TBK | 1:500 |  | Cell Signalling | 3504 |
| Anti-pTBK | 1:500 |  | Cell Signalling | 5483 |
| Anti-STING 1 | 1:500 |  | Cell Signalling | 13647 |
| Anti-pSTING | 1:500 |  | Cell Signalling | 50907 |
| Anti-CSA | 1:500 |  | Abcam | ab137033 |
| Anti-lamin A/C | 1:1000 |  | Santa Cruz | sc-7292 |
| Anti-Histone H3 | 1:1000 |  | Cell signalling | 3638 |
| Anti-GFP | 1:1000 |  | ThermoFisher | MA5-15256 |
| Anti-BAF | 1:250 |  | ProSci | 4019 |
| Anti-CSB | 1:500 |  | Abcam | ab96089 |
| Anti-DDB1 | 1:250 |  | BD Biosciences | 612488 |
| Anti-GAPDH | 1:5000 |  | Invitrogen | MA5-15738 |
| Anti-lamin B1 |  | 1:500 | Santa Cruz | sc-365214 |
| Anti-lamin A/C |  | 1:1000 | Santa Cruz | sc-376248 |
| Anti-emerin |  | 1:400 | Cell Signalling | 30853 |
| Anti-Vimentin |  | 1:100 | Cell signalling | 5741 |
| CF488A-phalloidin |  | 1:100 | Biotum | 00042-T |

| Secondary Antibodies |  |  |  |  |
| --- | --- | --- | --- | --- |
| Antibody | Western Blot Dilution | Immunofluorescence Dilution | Supplier | Catalogue Number |
| IRDye 800RD anti-mouse | 1:10000 |  | LI-COR | 925-32212 |
| IRDye 600 RD anti-rabbit | 1:10000 |  | LI-COR | 925-68073 |
| Alexa Fluor 647 anti-rabbit |  | 1:1000 | Life Technologies | A31573 |
| Alexa Fluor 488 anti-mouse |  | 1:1000 | Life Technologies | A21141 |
| Alexa Fluor 568 anti-mouse |  | 1:1000 | Life Technologies | A21124 |
| Alexa Fluor 488 anti-rabbit |  | 1:1000 | Life Technologies | A21206 |
| Alexa Fluor 647 anti-mouse |  | 1:1000 | Life technologies | A21242 |
| Alexa Fluor 488 anti-mouse |  | 1:1000 | Life technologies | A21121 |
| Alexa Fluor 568 anti-rabbit |  | 1:1000 | Life technologies | A10042 |
| Alexa fluor 647 anti-mouse |  | 1:1000 | Life technologies | A31571 |

### Immunoprecipitation (IP)

Cells were lysed in IP lysis buffer (0.5% NP-40, 150 mM NaCl, 0.5 mM EDTA, 10 mM Tris-HCl, pH = 7.4)) with freshly prepared 0.5 mM PMSF and cOmplete^TM^ Mini EDTA-free Protease Inhibitor Cocktail (Roche, #04693159001) on ice for 30 minutes with gentle vortexing every 10 minutes. Lysates were subjected to centrifugation at 15,000 g for 15 minutes at 4°C and the supernatant was collected. The remaining pellet was then resuspended in RIPA buffer freshly supplemented with 0.5 mM PMSF and cOmplete^TM^ Mini EDTA-free Protease Inhibitor Cocktail and subjected to sonification two times at 10 kHz for 10 seconds. The resuspended pellet solution was then centrifuged at 15,000 g for 15 minutes at 4°C and the supernatant was combined with the supernatant obtained from the first lysis. Input samples were collected from the combined lysates. For the pulldowns, ChemTek GFP-Trap Magnetic Agarose (Proteintech, #gtma) or ANTI-FLAG^®^ M2 Affinity Gel (Sigma-Aldrich, #A2220) were incubated with the protein lysates for 4 hours under rotation at 4°C. Protein-bound beads were then washed four times with wash buffer. Protein elution from the protein-bound beads was performed by adding Protein Sample Loading Buffer (Li-cor, #928-40004) with 10 mM DTT followed by boiling for 5 min at 95°C.

### Proximity Ligation Assay (PLA)

Cells were first seeded onto 12 mm coverslips and fixed in 4% (v/v) paraformaldehyde dissolved in phosphate-buffered saline (PBS) for 10 minutes at room temperature. Permeabilisation was performed by incubating cells with 0.2% (v/v) Triton X-100 dissolved in PBS for 15 minutes at room temperature. The following steps including blocking, primary antibody incubation, DuoLink Probe incubation, ligation, and amplification were performed using the DUOLink PLA assay kit (#DUO92008, Sigma Aldrich) following the manufacturer’s instructions. Primary antibodies for incubation were mouse anti-lamin A/C (Santa Cruz, #sc-376248, 1:1000) and rabbit anti-LEMD2 (Sigma Aldrich, #HPA017340, 1:250). For the Duolink In Situ PLA probe, anti-mouse MINUS and anti-rabbit PLUS were used. For amplification, Duolink Amplification red was used. Coverslips were then mounted using Duolink Mounting Media containing DAPI. Acquisition of microscopy images was performed with the Stellaris 5 confocal laser scanning microscope using the 40X oil immersion objective lens (Leica HC PLA APO, 1.3 NA).

### Immunofluorescence assay

For samples requiring pre-extraction of soluble proteins prior to immunofluorescence assay, cells were incubated with cytoskeletal buffer (CSK) (100 mM NaCl, 300 mM sucrose, 1 mM EGTA, 1 mM MgCl_2_, 1 mM DTT, 10 mM PIPES/KOH, 6.8 pH) for 5 minutes on ice. Cells were fixed in 4% (v/v) paraformaldehyde dissolved in phosphate-buffered saline (PBS) for 10 minutes at room temperature. Permeabilisation was performed with 0.2% (v/v) Triton X-100 in PBS for 15 minutes at room temperature. Blocking was done with 2% (v/v) BSA dissolved in PBS for 30 minutes at room temperature. Coverslips were then incubated with primary antibodies and secondary antibodies, both for 1 hour at room temperature. Details of primary antibodies and secondary antibodies are outlined in **Table 2**. Coverslips were mounted onto glass slides using Prolong Gold Antifade Reagent (Cell Signalling, #9071). Acquisition of microscopy images was performed with the Stellaris 5 confocal laser scanning microscope using the 40X oil immersion objective lens (Leica HC PLA APO, 1.3 NA).

### Automated CellProfiler^TM^ workflow

CellProfiler^TM^ V4.1.3 software was used to analyze images obtained from immunofluorescence assay to NE phenotypes, protein localization, and protein fluorescent signal intensity. All Cell Profiler pipelines used in this work can be made available upon request.

To assess nuclear morphology, the nuclear form factor is calculated for each identified nuclei to determine its roundness. The nuclear form factor is calculated by 4πA/P^2^, where A is the area of nucleus, and P is the perimeter of the nucleus. A perfectly round nucleus will have a form factor of 1 and a more deformed nucleus will have a form factor lower than 1. DAPI staining was used to identify individual nuclei using the ‘Identify Primary Objects’ setting, and border objects are excluded. Following the identification of nuclei, the nuclear form factor was calculated using the ‘Measure Object Size and Shape’ setting. To measure the LEMD2 intensity at the nucleus, the measurement was performed by measuring the signal intensity of the LEMD2 channel in the identified nucleus, defined by the DAPI mask. To identify the cGAS foci at the nucleus, the signal intensity of cGAS foci was enhanced relative to the rest of the image using the ‘Enhance Or Suppress Features’ setting. cGAS foci were then identified using the ‘Identify Primary Objects’ setting.

The pipeline used to identify nuclear bleb was a modified version of the pipeline described before(36). DAPI staining and lamin B1 staining were used to identify individual nuclei using the ‘Identify Primary Objects’ setting, and border objects are excluded. Blebs were identified by subtracting the pixel areas of DAPI and lamin B1 staining using the ‘Identify Tertiary Objects’ setting. Identified blebs were then filtered by minimum pixel area to remove small identified false blebs and edge pixels of the nuclei.

To quantify PLA foci, individual nuclei were first identified using ‘Identify Primary Objects’ setting, and border objects are excluded. Signal intensity of PLA foci was enhanced relative to the rest of the image using the ‘Enhance Or Suppress Features’ setting before the PLA foci were identified using the ‘Identify Primary Objects’ setting.

### RNA sequencing (RNA-seq) and differentially expressed (DE) gene analysis

The read count matrix of WT(HA-CSA) and CS-A were downloaded from the Gene Expression Omnibus (GEO) website (GSE87540) (37). The datasets were obtained from WT(HA-CSA) and CS-A cells grown in DMEM/HamF10 media with 10% (v/v) fetal calf serum and in 40 µg/mL gentamycin, without UV irradiation.

The Deseq2 R package (v.1.24.0) was used to analyse the publicly available RNA-seq data unless otherwise stated. The median of ratios method was used to normalise the raw count data using the ‘estamateSizeFactors’ function. To compare the gene expression level between CS-A and WT, log2 fold change and p-values were calculated using the ‘nbinomWaldTest’ function. The p-values obtained from the Wald tests were adjusted for multiple testing using the Benjamini-Hochberg method. The list of significantly upregulated and downregulated genes was determined by subsetting DE genes with adjusted p-values less than 0.001 and absolute value of log2 fold change greater than 2. Gene ontology (GO) analysis and STRING protein-protein interaction diagram of the DE genes were perform using the STRING website (<u>STRING: functional protein association networks</u> (string-db.org)). The volcano plot was generated using the ‘EnhancedVolcano’ function from the EnhancedVolcano R Package to show significant DE genes between CS-A and WT. The adjusted p-values and log2 fold change cut-off were the same as per subsetting significantly upregulated and downregulated genes.

### Fluorescence recovery after photobleaching (FRAP)

WT(HA-CSA) and CS-A cells transiently expressing LEMD2-GFP fusion protein were imaged using 60X water-immersion objective lens (Zeiss, 1.2NA) Zeiss 880 Airyscan Confocal microscope. Cells were seeded onto a µ-slide 4 well chambered coverslip (ibidi, #80426) in CO_2_ independent medium (Gibco, #11580536) containing 10% (v/v) FBS and imaged at 37°C using the microscope cage incubator. The fluorescence intensity of the ROI was measured over 93 seconds at 1 second interval (93 images in total) using 2% laser power from 488 nm light. Photobleaching was started after 3 scans and recovery was followed by 90 scans. For photobleaching, 2.52 x 6.30 µm region of LEMD2-GFP fluorescence at the nuclear periphery in the mid-focal plane was photobleached by scanning 50 iterations using 100% light intensity from 488 nm light. The pinhole size was set at 1 AU for the confocal.

Analysis of FRAP data was done using Microscoft Excel Version 16.43 and GraphPad Prism Version 9. Fluorescence intensity measurement of ROI was corrected by the fluorescence intensity of an unbleached area at the nuclear periphery to account for background bleaching. The data was fitted with a curve of the form *y* = *y*_0_ + (*a* – *y*_0_)(1 – *e*^−*bk*^), where (*a*, *b*) corresponds to the asymptotic values of relative LEMD2-GFP fluorescence intensity and the decay rate of growth, respectively. The percentage of immobile fraction (*IF*%) was determined by *IF*% = *a* ⋅ 100% and half-time of recovery (*t*_1/2_) was determined by 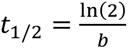.

### Live cell proliferation assay

WT(HA-CSA) and CS-A cells were seeded onto 24-well plate at ∼35% confluency and transfected with either the control siRNA or SUN1-targeting siRNA. To induce bulky DNA adducts mimicking the ones caused by UV irradiation, the cells were treated with 1.5 µM of 4-nitroquinoline 1-oxide (4NQO) for 1 hour prior to imaging. Phase contrast images were acquired every 4 hours over a period of 48 hours using an Incucyte machine (Sartorius). The percentage of cell confluence was calculated using the attached Incucyte^®^software (Sartorius).

### Statistical Analysis

All statistical tests and graphs were generated using GraphPad Prism Version 9. Error bars in graphs are shown as mean ± SD. Post-hoc tests were performed for experiments that required correction for multiple comparisons. Details of specific statistical tests are described in the figure legends.

## Results

NE defects are characteristic of premature ageing conditions associated with mutations in NE proteins, such as Hutchinson-Gilford progeria syndrome (HGPS), restrictive dermopathy, atypical progeria syndrome (APS), and Nestor-Guillermo progeria syndrome (NGPS) (3). In addition, there is now mounting evidence that NE dysfunction can also occur in physiological ageing (38). To further investigate the connection between NE dysfunction and ageing phenotypes, we decided to assess NE integrity in Cockayne Syndrome, a premature ageing condition that is not caused by NE-associated mutations. Interestingly, we observed that knocking out CSA but not CSB in HAP1 cells induced aberrant nuclear morphology (**Figure 1A**). To characterize the NE phenotypes further, we obtained CS patient-derived cell lines carrying loss-of-function mutations in CSA (CS-A cells) or CSB (CS-B cells), and their respective isogenic control cell lines (WT(HA-CSA) and WT(HA-GFP-CSB)). The expression of HA-CSA and HA-GFP-CSB in the isogenic control cell lines and the absence of CSA and CSB expression in CS-A and CS-B cells, respectively, were confirmed by immunoblotting **(Figure 1B)**. We first assessed the nuclear shape and the presence of nuclear blebbing. Nuclear blebs are a good proxy to assess NE ruptures as blebs occur where the NE is weakened, and almost always result in NE ruptures (30,36). CS-A cells showed misshapen nuclei with a significantly lower nuclear circularity (represented by the reduced form factor) and a higher percentage of blebbing compared to the WT(HA-CSA) **(Figure 1C, 1D)**. On the contrary, CS-B cells showed a slight increase of nuclear circularity and no difference in the percentage of blebbing compared to WT(HA-GFP-CSB) cells **(Figure 1E, 1F)**. As a more direct indication of NE rupture, we quantified the percentage of nuclei with cyclic GMP-AMP synthase (cGAS) foci. cGAS is a dsDNA sensor that binds to cytosolic DNA, and upon NE rupture, cGAS binds to the genomic DNA being exposed at the site of NE rupture (39). Consistent with the increased blebbing, we observed that CS-A cells display more cGAS foci per nucleus compared to the WT(HA-CSA) **(Figure 1G, 1H)**, whereas no significant differences were detected between CS-B and WT(HA-GFP-CSB) cells **(Figure 1I, 1J)**. Together, these observations show the presence of NE deformation, blebbing, and ruptures upon CSA loss of function mutation in patient cells.

**Figure 1:**
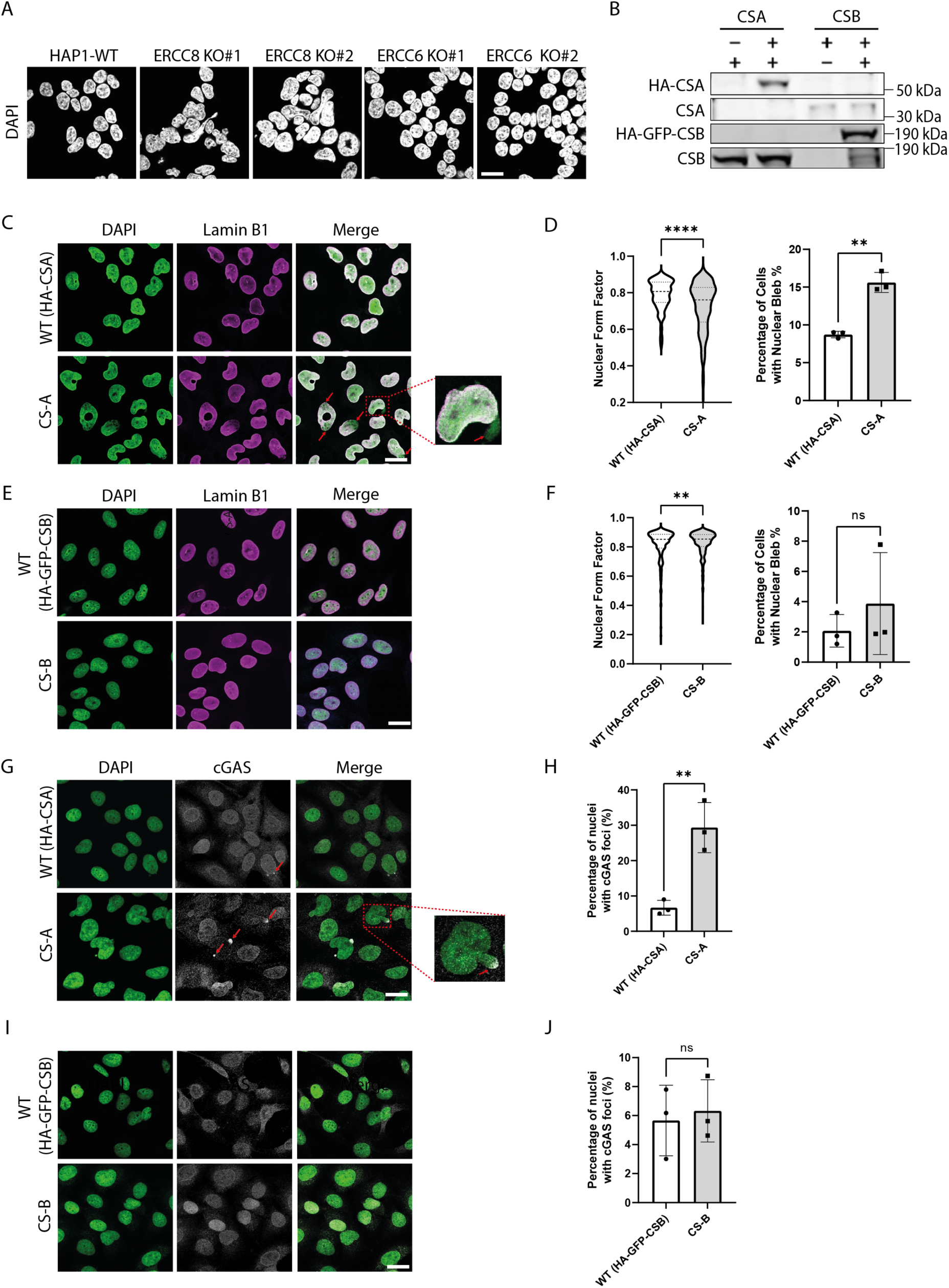
Loss of CSA but not CSB induces NE defects. **(A)** Representative DAPI staining images showing the effect of CSA or CSB knockout on nuclear shape in HAP1 cell line (representative images of two independent CRISPR-knockout clones). **(B)** Immunoblot showing the expression of CSA and CSB in the indicated cell lines. **(C)** Representative DAPI and lamin B1 immunofluorescence images of WT(HA-CSA) and CS-A cells; scale bar: 25 µm. Red arrows indicate nuclear blebs and zoom in image shows an example of nuclear bleb. **(D)** Quantification of nuclear form factor and percentage of blebbing for WT(HA-CSA) and CS-A cells, respectively, using two-tailed unpaired t-test (** p < 0.01, **** p < 0.0001), in n=3 with >300 cells per experiment. **(E)** Representative DAPI and lamin B1 immunofluorescence images of WT(HA-GFP-CSB) and CS-B cells; scale bar: 25 µm. **(F)** Quantification of nuclear form factor and percentage of blebbing for WT(HA-GFP-CSB) and CS-B cells, respectively, using two-tailed unpaired t-test (ns p > 0.05, ** p < 0.01), n=3 with >300 cells per experiment. **(G)** Representative DAPI and cGAS immunofluorescence images of WT(HA-CSA) and CS-A cells; scale bar: 25 µm. Red arrows indicate cGAS foci and zoom in image shows an example of cGAS foci. **(H)** Quantification of the percentage of nuclei with cGAS foci for WT(HA-CSA) and CS-A cells, respectively, using two-tailed unpaired t-test (** p < 0.01), n=3 with >300 cells per experiment. **(I)** Representative DAPI and cGAS immunofluorescence images of WT(HA-GFP-CSB) and CS-B cells; scale bar: 25 µm. **(J)** Quantification of the percentage of nuclei with cGAS foci for WT(HA-GFP-CSB) and CS-B cells, respectively, using two-tailed unpaired t-test (ns p > 0.05), n=3 with >300 cells per experiment. All experiments in this figure were n=3 independent experiments.

To explore whether the loss of CSA may cause NE defects by affecting the level of NE proteins, we probed a panel of “core” NE proteins including lamin A, lamin C, lamin B1, LEMD2, emerin, and SUN1 by Western blotting **(Figure 2A)**. There was no noticeable difference in the expression of these proteins between WT(HA-CSA) and CS-A cells. We then assessed the subcellular localization of these NE proteins using immunofluorescence. Similarly, there was no noticeable difference between WT(HA-CSA) and CS-A cells **(Figure 2B)**. For LEMD2, due to high background signal given by the antibody, it was challenging to visualize its NE-localisation. Therefore, we performed pre-extraction to visualize the ‘insoluble’ pool of LEMD2 **(Figure 2C)**. The specificity of the detected LEMD2 signal post pre-extraction was confirmed using a LEMD2 siRNA **(Figure S1)**. We showed that the amount of insoluble LEMD2 was significantly lower (∼20%) in CS-A cells compared to WT(HA-CSA) **(Figure 2D)**. This result was confirmed by western blotting following a similar pre-extraction, while the LEMD2 expression level in the total cell lysate was unchanged **(Figure 2E, 2F)**.

**Figure 2:**
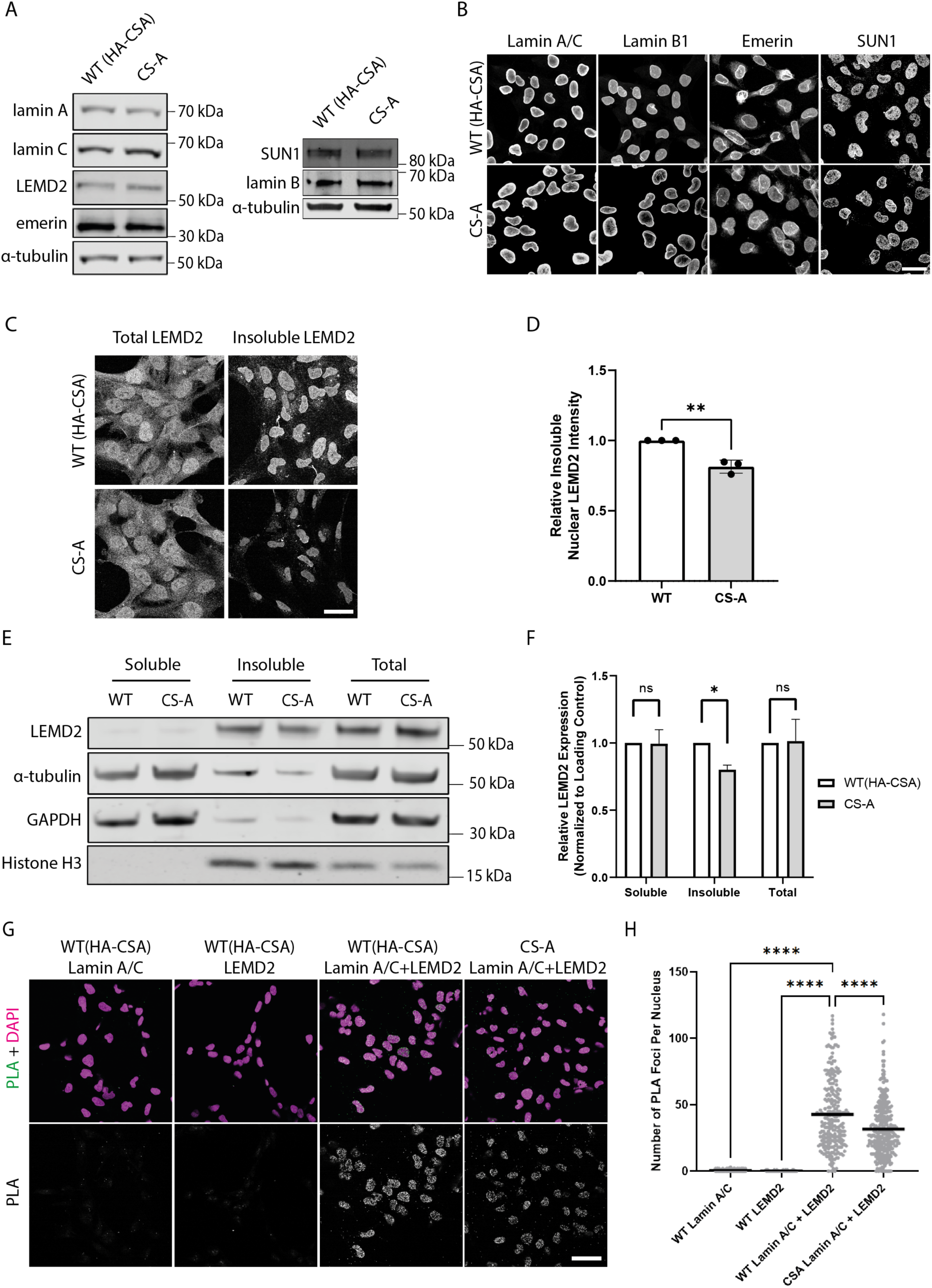
LEMD2 incorporation into the NE is decreased in CS-A cells: **(A)** Representative Western blot of the indicated NE proteins in WT(HA-CSA) and CS-A cells. **(B)** Representative immunofluorescence images of lamin A/C, lamin B1, emerin, and SUN1 staining; scale bar: 25 µm. **(C)** Representative immunofluorescence staining showing total and insoluble LEMD2 protein in WT(HA-CSA) and CS-A cells; scale bar: 25 µm. **(D)** Quantification of the insoluble LEMD2 showing a significant reduction in CS-A compared to WT(HA-CSA) using two-tailed unpaired t-test (** p < 0.01), n=3 with >100 cells per experiment. **(E)** Representative immunoblot showing LEMD2 expression level in the soluble, insoluble, and whole cell extracts in WT(HA-CSA) and CS-A cells. **(F)** Quantification of the relative LEMD2 expression level from the western blots as shown in (E); normalised to GAPDH, histone H3, and α-tubulin, respectively. P values were calculated using two-tailed paired t-tests (ns p > 0.05, * p < 0.05). **(G)** Representative confocal images of DAPI and PLA signal in the indicated cells using either anti-lamin A/C or anti-LEMD2 antibodies alone (negative controls) or in combination; scale bar: 50 µm. **(H)** Quantification of the number of PLA foci per nuclei. P value was calculated using one-way ANOVA test followed by Tukey’s post hoc test (**** p < 0.0001), n=3 with >100 cells per experiment. All experiments in this figure were n=3 independent experiments.

LEMD2 is a known A-type lamin-binding protein and previous literature suggests that the binding of these two proteins is important for maintaining the structural integrity of the nucleus (40). Since our data suggested a decrease of LEMD2 present at the NE in CS-A cells, we speculated that this could lead to decreased LEMD2 binding to lamin A/C at the NE which may cause NE fragility. Using a Proximity Ligation Assay (PLA), we showed a significant reduction in the number of PLA foci in CS-A cells compared to the WT(HA-CSA) cells, reflecting a reduced number of LEMD2-lamin A/C complexes **(Figure 2G, 2H)**. This data suggests defects in the incorporation of LEMD2 into the NE and lamin protein complexes in CS-A cells.

To understand the potential mechanism by which the NE incorporation of LEMD2 might be affected in CSA-depleted cells, we then investigated a potential interaction between LEMD2 and CSA. Using BioID approaches, previous literature demonstrated that LEMD2 is in proximity with known interactors of CSA, including CUL4A, DDB1, and CSN (41). Therefore, we hypothesized that LEMD2 could also interact with CSA and this interaction may be important to properly incorporate LEMD2 into the NE. To address this, we overexpressed LEMD2-GFP and Flag-CSA constructs, followed by GFP pulldown in WT(HA-CSA) cells. We observed that LEMD2-GFP co-immunoprecipitated with HA-CSA and Flag-CSA **(Figure 3A)**. BAF was used as a positive control for known LEMD2 interactors (31,35). This finding suggests that CSA can indeed interact with LEMD2.

**Figure 3:**
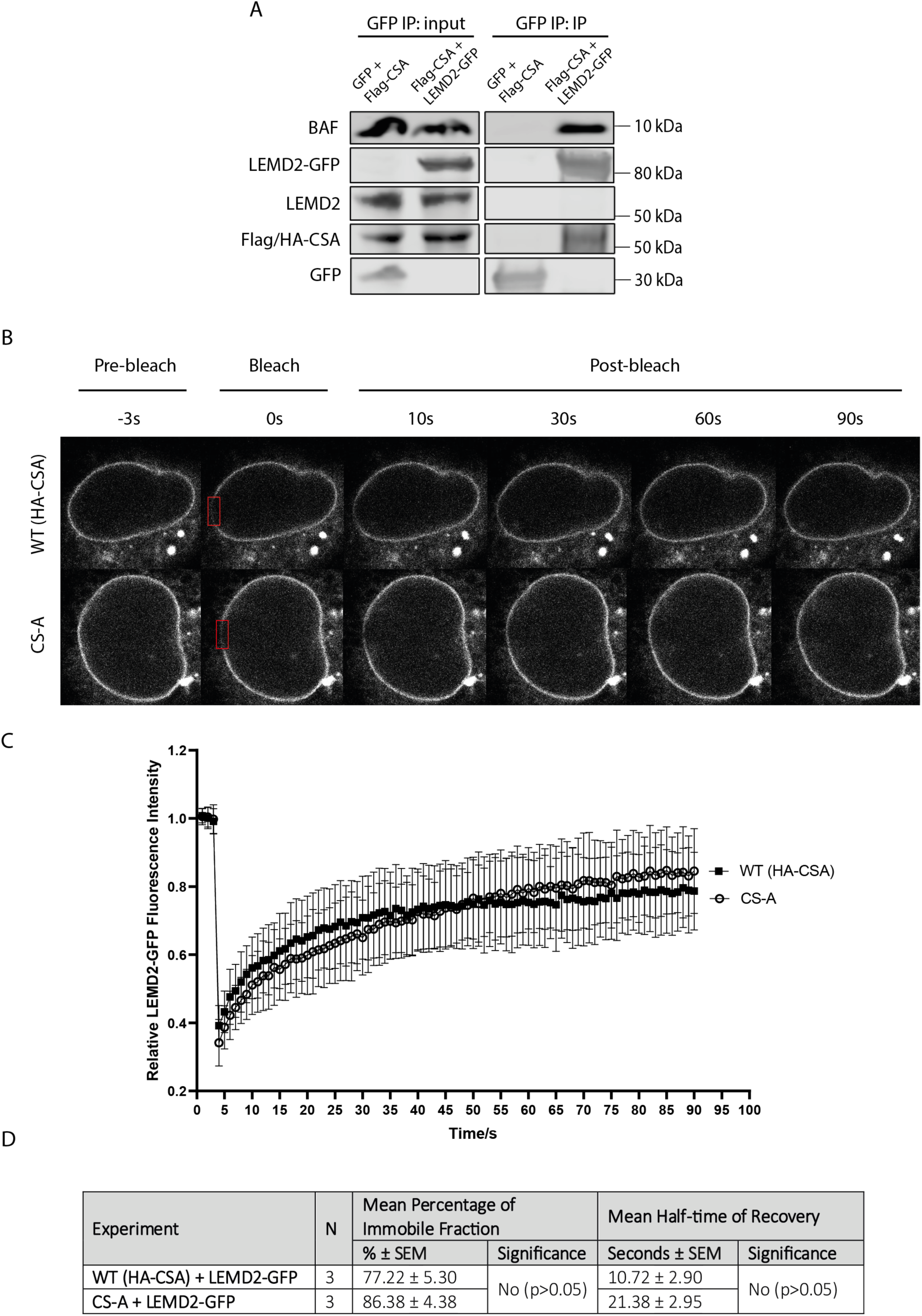
The absence of CSA does not affect LEMD2 mobility at the NE: **(A)** Immunoprecipitation of LEMD2-GFP in WT(HA-CSA) cells overexpressing GFP+Flag-CSA (control) and Flag-CSA+LEMD2-GFP. **(B)** Representative time-lapse confocal images showing pre-bleach, bleach, and post-bleach images of the WT(HA-CSA) and CS-A nuclei with LEMD2-GFP overexpression. The red box indicates the photobleached region at the nuclear periphery. **(C)** Graph showing the FRAP kinetics of LEMD2-GFP expressed in WT(HA-CSA) (n=25 cells) and CS-A (n=25 cells) cells. Error bars represent standard deviation. **(D)** Summary table showing the mean percentage of the immobile fraction and the mean half-time of recovery of LEMD2-GFP FRAP experiment. Data were compared using two-tailed unpaired t-test. All experiments in this figure were n=3 independent experiments.

To establish whether this interaction may be important in stabilizing or immobilizing LEMD2 at the NE, we performed FRAP experiments in WT(HA-CSA) and CS-A cells transiently expressing LEMD2-GFP **(Figure 3B)**. Both WT(HA-CSA) and CS-A cells showed similar LEMD2-GFP recovery half-time and percentage of immobile LEMD2-GFP fraction **(Figure 3C, 3D)**. Altogether, this data suggests that CSA can bind LEMD2 and that the absence of CSA in CS-A patient cells does not affect the mobility of LEMD2 at the NE but instead decreases its interaction with A-type lamins.

In searching for other mechanisms that could contribute to the loss of NE integrity in CS-A cells, we analyzed the publicly available bulk RNA-seq dataset (GSE87540) obtained from WT(HA-CSA) and CS-A cells (37). Through DE gene analysis, we identified genes that were significantly upregulated and downregulated in CS-A cells compared to WT(HA-CSA). Through gene ontology analysis, we found that genes involved in endoplasmic reticulum (ER) stress were differentially expressed **(Figure 4B)**, which could be a consequence of the loss of proteostasis in CS-A cells as described previously (42,43). More surprisingly, we also identified differential transcript expressions of genes involved in cytoskeleton polymerization, including F-actin, alpha-tubulin, and vimentin **(Figure 4A, 4B)**. Since the cytoskeleton has a well-established role in regulating nuclear mechanical properties through binding to the LINC complex (44), we analysed F-actin, alpha-tubulin, and vimentin by immunofluorescence in CS-A cells **(Figure 4C)**. There were no obvious changes in the alpha-tubulin and the vimentin network organisation between the WT(HA-CSA) and CS-A cells. However, we observed a clear increase in the formation of actin stress fibers in CS-A cells compared to WT(HA-CSA). Actin stress fibers are contractile molecular bundles in non-muscle cells that consist of parallel actin and myosin II filaments, and other actin-binding proteins including filamins, fascins, and actinins (45). We therefore decided to further pursue how mechanical forces generated by actin stress fibers may contribute to the loss of NE integrity in CS-A cells.

**Figure 4:**
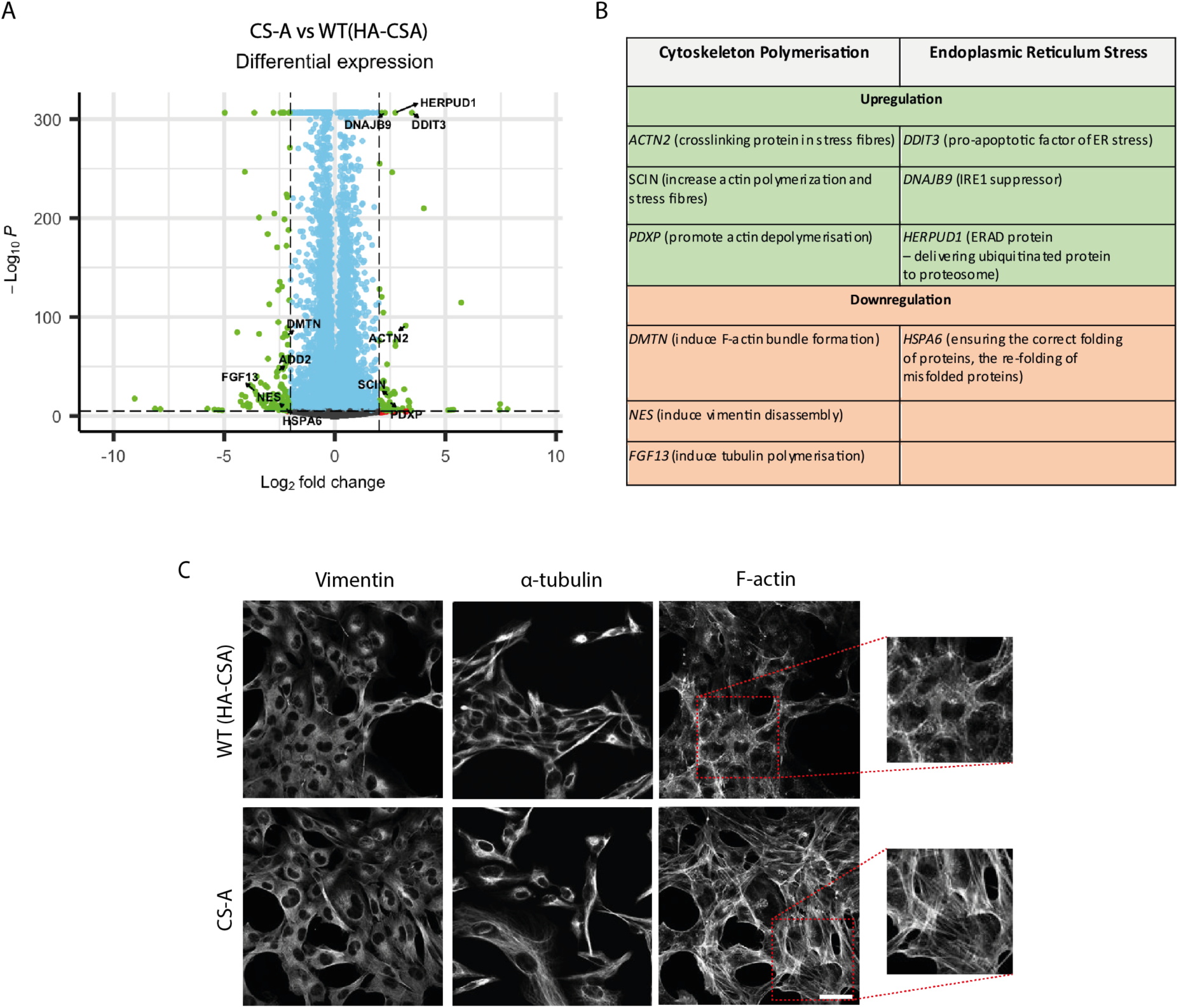
Loss of CSA results in transcriptional deregulation of cytoskeleton polymerization-regulating genes. **(A)** Volcano plot showing differentially expressed genes in CS-A and WT(HA-CSA) cells. (**B**) Table showing significantly upregulated and downregulated genes involved in cytoskeleton polymerization and endoplasmic reticulum stress. **(C)** Representative immunofluorescence staining showing the appearance of F-actin stress fibers in CS-A cells compared to WT(HA-CSA), but no noticeable difference in α-Tubulin, and vimentin networks; scale bar: 50 µm. All experiments in this figure were n=3 independent experiments.

To this aim, we disrupted actin polymerization using Cytochalasin D, a potent inhibitor of actin polymerisation **(Figure 5A)**. Interestingly, Cytochalasin D treatment significantly improved nuclear circularity and decreased the number of nuclear blebs in CS-A cells, indicating an improvement in the NE integrity **(Figure 5B, 5C, 5D)**. We then sought to perform a reverse experiment by treating WT(HA-CSA) and CS-A cells with Jasplakinolide **(Figure 5A)**, which is an actin polymerization-inducing drug that stimulates the nucleation of actin filaments. We hypothesized that by stabilizing actin stress fibers in WT(HA-CSA) cells with Jasplakinolide, the Jasplakinolide-treated WT(HA-CSA) cells would display NE defects similar to what we observed in untreated CS-A cells. Consistent with our hypothesis, WT(HA-CSA) cells treated with Jasplakinolide displayed a significant reduction in nuclear circularity and increased nuclear blebbing **(Figure 5B, 5C, 5D)**. When treating CS-A cells with Jasplakinolide, no significant change in either the nuclear circularity or percentage of nuclear blebs was detected. This suggests that enhancing actin polymerization or stabilization cannot further increase the NE defects already present in CS-A cells. All together, these data suggest that the transcriptional deregulation of genes affecting actin polymerization in CS-A cells contributes to the loss of NE integrity.

**Figure 5:**
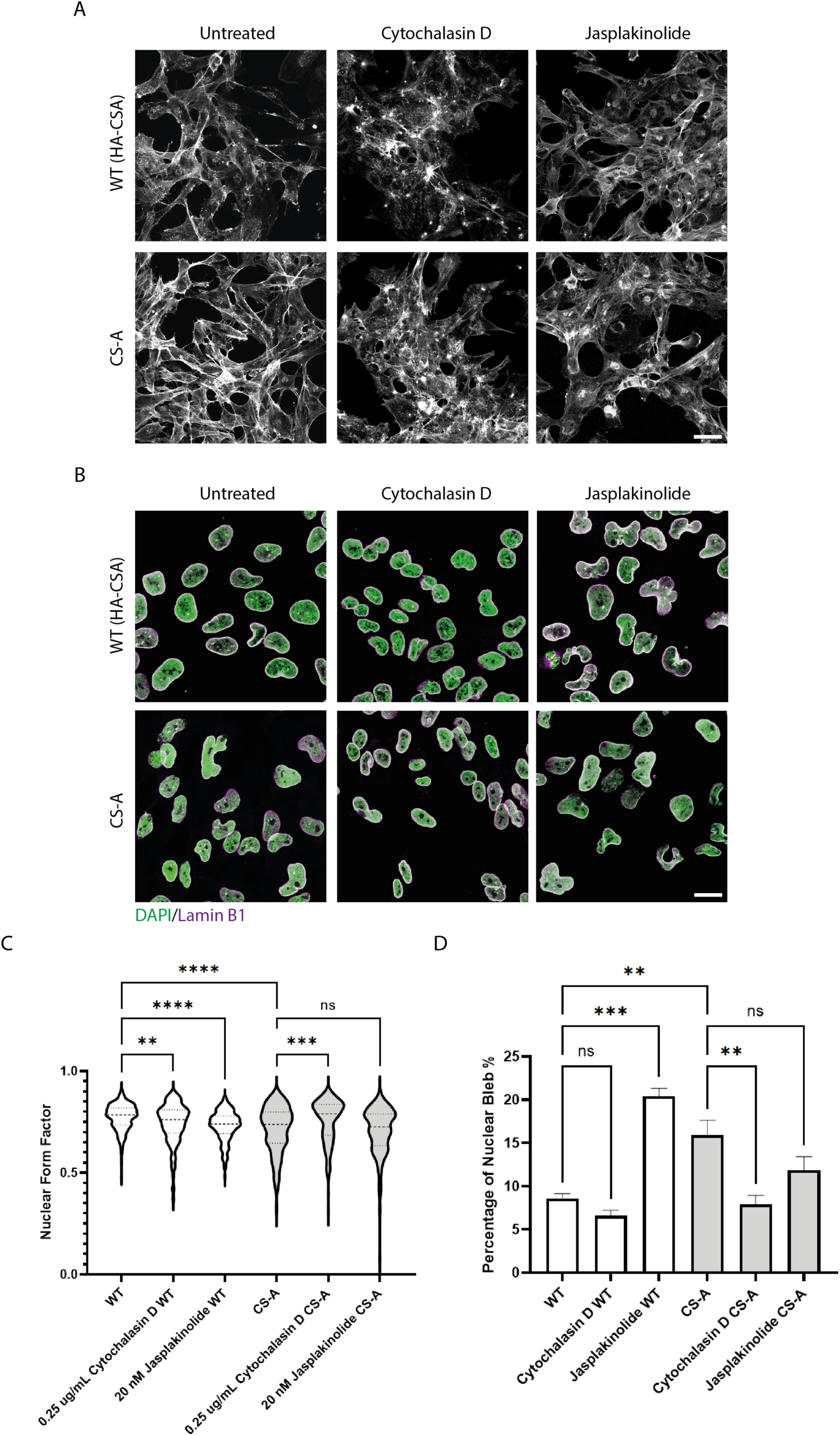
Modulation of actin polymerization affects the NE phenotypes in CS-A cells. **(A)** Immunofluorescence staining of F-actin in WT(HA-CSA) and CS-A cells treated with 0.25 µg/mL Cytochalasin D and 25 nM Jasplakinolide for 24 hours; scale bar: 50 µm. **(B)** Representative immunofluorescence staining of DAPI and lamin B1 in WT(HA-CSA) and CS-A cells treated with the actin polymerization inhibitor (0.25 µg/mL Latrunculin A) or the actin stabilizer (25 nM Jasplakinolide) for 24 hours; scale bar: 25 µm. **(C)** Quantification of the nuclear form factor and **(D)** percentage of nuclear blebs in Cytochalasin D treated cells, n=3 with >100 cells per experiment. P value was calculated using one-way ANOVA test followed by Tukey’s post hoc test (ns p > 0.05, ** p < 0.01, *** p < 0.001, **** p < 0.0001). All experiments in this figure were n=3 independent experiments.

Cytoskeleton stress fibers can transduce mechanical forces to the NE through the LINC complex, consisting of the KASH-domain and SUN-domain proteins (25,44). Therefore, we asked whether disrupting the cytoskeletal-to-NE interaction in CS-A cells by depleting components of the LINC complex could relieve the mechanical forces at the NE generated by actin stress fibers and improve the NE phenotypes. Previous studies showed that SUN1, but not SUN2, is the main interactor of KASH-domain proteins in the LINC complex assembly, and depleting SUN1 inhibits NE rupture in cancer cell lines (46). For that reason, we knocked down SUN1 using siRNA in WT(HA-CSA) and CS-A cells **(Figure 6A)**. Immunofluorescence staining of DAPI and lamin B1 was then performed in siSUN1-depleted WT(HA-CSA) and CS-A cells **(Figure 6B)** to identify nuclear blebs (36,47). We observed that the depletion of SUN1 in CS-A cells increased nuclear circularity and significantly reduced the number of nuclear blebs **(Figure 6C, 6D)**. This finding suggests that the LINC complex mediates NE deformation and rupture in CS-A cells, possibly through transmitting increased mechanical forces generated by actin stress fibers to the NE.

**Figure 6:**
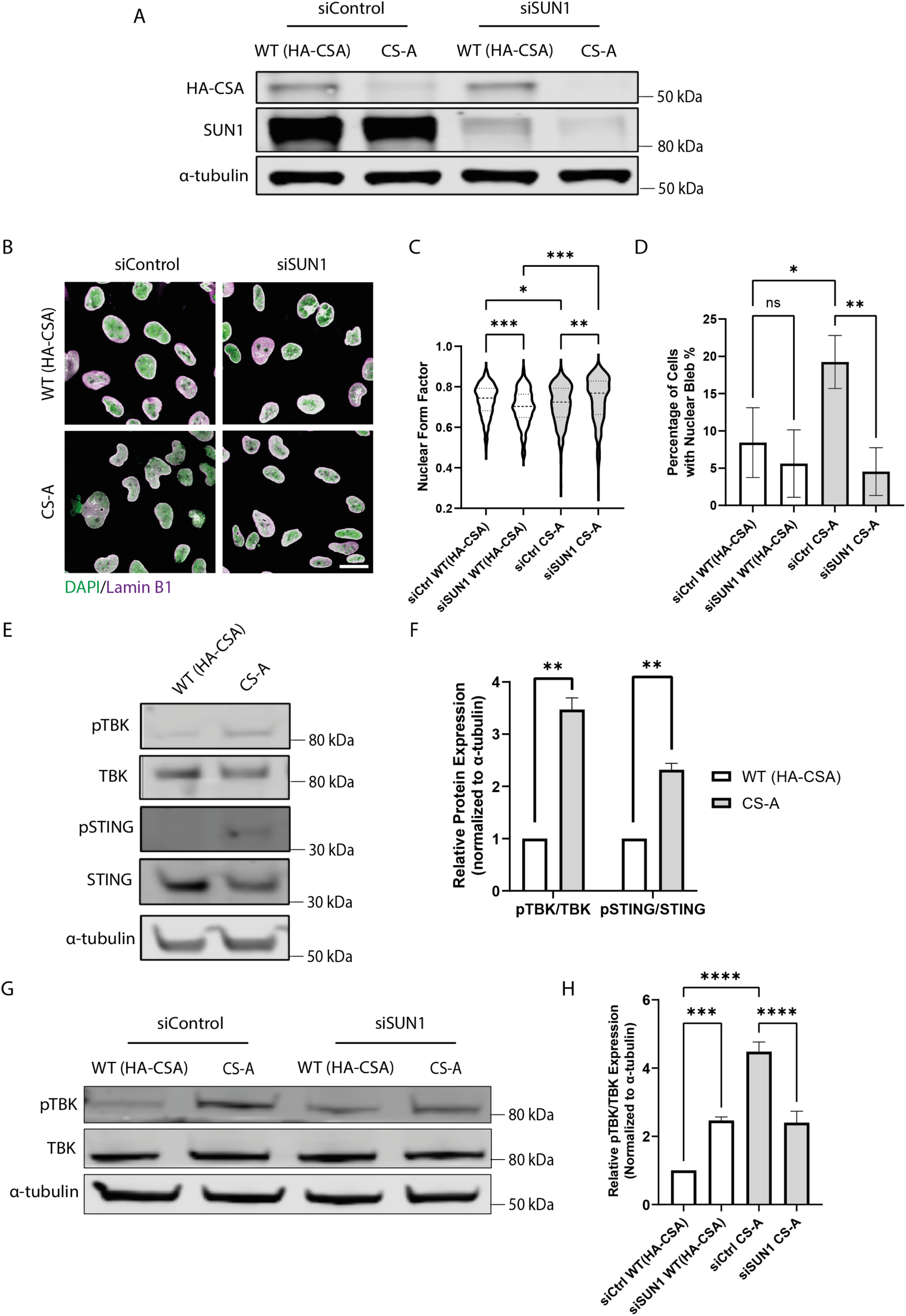
Disruption of the LINC complex improves NE phenotypes in CSA-depleted cells. **(A)** Immunoblot showing the efficiency of SUN1 knockdown by siRNA in WT(HA-CSA) and CS-A cells. **(B)** Representative immunofluorescence staining of DAPI and lamin B1 in WT(HA-CSA) and CS-A cells transfected with SUN1-targeting siRNA (siSUN1**). (C)** Quantification of the nuclear form factor and **(D)** percentage of nuclear blebs, n=3 with >100 cells per experiment. siCtrl: scramble siRNA, siSUN1: siRNA targeting SUN1. P value was calculated using one-way ANOVA test followed by Tukey’s post hoc test (ns p > 0.05, * p < 0.05, ** p < 0.01, *** p < 0.001). **(E)** Representative western blot showing phosphorylation levels of TBK and STING in WT(HA-CSA) and CS-A cells. **(F)** Quantification of the relative phosphorylation level of cGAS-STING pathway proteins. P value was calculated using paired t-test (** p < 0.01). **(G)** Representative western blot showing phosphorylation levels of TBK in WT(HA-CSA) and CS-A cells upon SUN1 depletion. **(H)** Quantification of the relative phosphorylation level of TBK in SUN1-depleted cells P value was calculated using one-way ANOVA test followed by Tukey’s post hoc test (*** p < 0.001, **** p < 0.0001). All experiments in this figure were n=3 independent experiments.

As CS-A cells showed accumulation of cGAS foci **(Figure 1G, 1H)**, we assessed whether this was associated with the activation of the cGAS/STING pathway upon NE rupture. Upon cGAS binding to DNA, cGAS can activate and trigger the phosphorylation of downstream effectors including TBK and STING (32,48). Indeed, we observed increased phosphorylation levels of TBK and STING in CS-A cells **(Figure 6E, 6F)**. Preventing the NE stress in CS-A cells by depleting SUN1 was able to reduce the cGAS/STING pathway activation **(Figure 6G, 6H)**. Together, our results show that increased mechanical forces generated by the cytoskeleton contribute to NE defects in CS-A cells, causing ruptures that lead to the cGAS/STING pathway activation which can be ameliorated by LINC complex disruption.

One of the best-established characteristics of CS-A cells is their sensitivity to DNA damage generated by UV irradiation (5), due to the role of CSA in TC-NER. Therefore, we wondered whether the function of CSA in NE regulation was related in any way to its known function in DNA damage repair. To address this question, we set up a survival assay in response to the damage induced by the UV-mimetic chemical 4NQO. As expected, the loss of CSA resulted in a significant drop of cell survival after exposure to 4NQO, compared to the WT(HA-CSA) cells **(Figure 7A)**. Interestingly, SUN1 depletion was not able to improve the survival of CS-A cells in response to 4NQO **(Figure 7A-B).** This experiment shows that even when the NE defects and ruptures are rescued through SUN1 depletion, the CS-A cells are still sensitive to DNA damage. This result suggests that the function of CSA in maintaining the NE integrity is independent of its known function in TC-NER.

**Figure 7:**
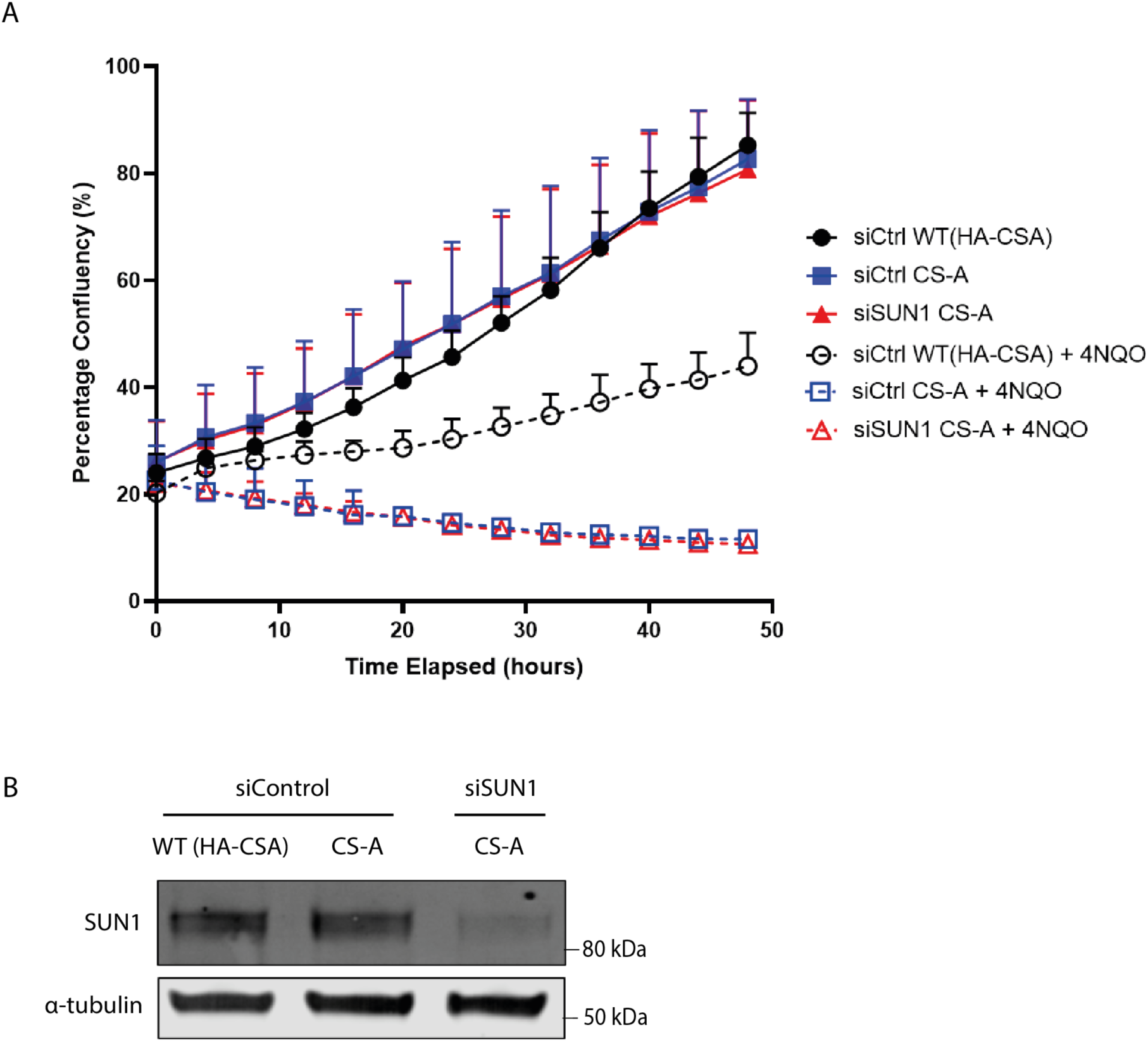
SUN1-depletion does not rescue the UV sensitivity phenotype of CS-A cells. **(A)** WT(HA-CSA) and CS-A cells transfected with control and SUN1 siRNA were treated for 1-hour with 4NQO, then imaged every 4 hours for 48 hours using an Incucyte system. Cell confluency was assessed with the Incucyte^®^ software. Error bars represent standard deviation. **(B**) Immunoblot showing the depletion of SUN1 in CS-A cells as presented in (A).

## Discussion

In this work, we report for the first time that loss of function of CSA in CS-A patient cells, leads to NE defects. Loss of NE integrity is a well-described feature of cells from HGPS patients and other laminopathies associated with NE dysfunction (49). This is reflected by the presence of nuclear deformation, NE ruptures, and lamin invaginations (50). More recently, NE defects have also been observed in cells from normally aged individuals, as well as in age-associated pathologies including neurodegeneration (51–53). Here, with the aim of exploring the potential contribution of NE dysfunction to other premature ageing pathologies, we observed loss of NE integrity in cells from Cockayne Syndrome A patients. These cells displayed a reduction in the NE-associated LEMD2 protein in CS-A cells, resulting in decreased formation of lamin A/C-LEMD2 complexes at the NE. LEMD2 has a well-established role in the maintenance of NE morphology and cell survival (40,54). Our findings are consistent with a previous study showing that LEMD2 depletion leads to NE defects without affecting the localization or expression of lamin A/C and lamin B1 (40). The authors speculated that although lamin proteins still localized to the NE, the lack of LEMD2 connected to the lamina network resulted in NE fragility. Since LEMD2 interaction withl lamin A/C is reduced in CS-A cells, this could contribute to the destabilization of the lamina, causing NE instability. The NE phenotypes observed in CS-A cells also bear similarities to those observed in Marbach-Rustad Progeria Syndrome (MRPS). MRPS is a recently characterized premature ageing disorder caused by a *de novo* mutation (c.1436C>T, p.S479F) in the C-terminal domain of LEMD2 (55). The mutation causes ’patchy’ localization of LEMD2 within the NE without affecting the total LEMD2 protein expression. Both MRPS and CS-A patient fibroblasts exhibit reduced nuclear circularity and increased nuclear blebbing phenotypes, further supporting the notion that the reduced LEMD2 at the NE may underlie the pathogenic NE phenotype in CS-A cells.

Mutations in LEMD2 have also been linked to cardiomyopathy in humans (56). For instance, patients carrying homozygous mutation (c.38T>G, p.L13R) in the *LEMD2* gene develop ventricular arrhythmia and fibrosis (57). It is noteworthy to mention that the L13R mutation causes a significant reduction in LEMD2 expression in cardiomyocytes and causes NE defects including nuclear membrane invaginations and decreased nuclear circularity. MRPS patients with a mutation in *LEMD2* also display cardiovascular defects with septal hypertrophy and right bundle branch block. However, CSA patients do not display any form of cardiomyopathy. A study in mice by Ross et al. (58) showed that reducing Lem2 levels in adult mice cardiomyocytes by approximately 45% did not lead to NE defects or cardiac dysfunction, suggesting a redundant function of LEMD2 at the NE in mouse heart. As our data show a reduction of LEMD2 of about 20% at the NE in CS-A cells compared to WT, it is maybe not surprising that the remaining LEMD2 at the NE is sufficient to maintain NE function in CSA patients’ cardiomyocytes.

Recent work suggested the LEMD2-specific interactome included CUL4A, DDB1, and CSN, which together form an E3 ubiquitination ligase complex with CSA (41). Here, we showed by immunoprecipitation that LEMD2 also interacts with CSA. This suggests that the recruitment and stabilization of LEMD2 to the NE is mediated by an interaction with CSA, although the mechanism remains unclear. Our FRAP experiment showed that the presence or absence of CSA did not influence LEMD2 mobility. The potential caveat here is that the GFP-LEMD2 overexpression, required to carry out this experiment, may have been enough to rescue a normal NE composition in CS-A cells. An alternative method to bypass this would be to engineer endogenous tagging of LEMD2 with GFP, to allow the study of LEMD2 protein motility without overexpression.

In our search for new molecular mechanisms contributing to NE defects in CS-A cells, and because previous literature suggested that CS was associated with broad transcriptional changes, we analyzed publicly available RNA-seq data from WT(HA-CSA) and CS-A cells obtained in absence of damage caused by UV irradiation (37). We found CS-A cells displayed a specific transcriptional deregulation of cytoskeletal proteins. Our data confirmed an increase in actin stress fibers in CS-A cells. The contribution of the actin cytoskeletal forces to spontaneous NE ruptures was established by a study conducted by Hatch and Hetzer (46). The authors showed that for cells growing on a flat and rigid 2D substrate (such as the ones used in standard culture conditions), contractile actin compresses the nucleus, leading to chromatin herniations and NE ruptures. Similarly, our findings show that disruption of actin polymerization was enough to improve the NE abnormalities in CS-A cells. On the other hand, chemically induced stabilization of the actin network in WT(HA-CSA) cells induced NE defects similar to that of the untreated CS-A cells. This data has reinforced the role of actin stress bundles in inducing spontaneous NE ruptures and has also highlighted how actin stress fibers in CS-A cells, occurring through transcriptional deregulation, contribute to the appearance of NE deformation and ruptures.

Another way to release mechanical stress on the nucleus is to disrupt the connection between actin and the NE by interfering with the integrity of the LINC complex. As such, depletion of SUN1 - a major component for LINC complex assembly - has been reported to correct several pathological phenotypes in HGPS and lamin A/C deficient cells including nuclear shape, NE blebbing, heterochromatin loss, chromatin disorganization, and cellular senescence (59). *In* vivo data also showed beneficial effect where removal of SUN1 in HGPS and lamin A/C deficient mouse models improved longevity and multiple pathological phenotypes including body weight deficit, lordokyphosis, trabecular, and bone densities (59). Similarly, SUN1 depletion led to the improvement of NE phenotypes in CS-A cells, reinforcing the idea that the forces exerted by actin on the nucleus of CS-A cells contribute to the NE abnormalities and ruptures in these cells. This data also suggests that LINC complex disruption could be an effective strategy to restore cellular homeostasis in multiple syndromes associated with NE dysfunction, and it is an approach that is currently being investigated by other teams (60,61).

As a result of the NE ruptures in CS-A cells, we observed the activation of the innate immune cGAS/STING pathway which was reduced upon SUN1 depletion in CS-A cells. Activation of cGAS/STING can induce the transcription factor NF-κB which in turn upregulates the release of proinflammatory chemokines, cytokines, and growth factors (48,62). These immune modulators are known as senescence-associated secretory phenotype (SASP) and act in an autocrine and paracrine manner to induce propagation and amplification of senescence in distant cells (63). In many diseases, SASP is the main contributor to chronic inflammation and progression of fibrosis (64,65). The only *in vivo* study suggesting inflammatory phenotype in CS model was done using CSA knockout (referred to as CX) mice (66). They observed increased senescence of brain endothelial cells and an upregulation of proinflammatory markers in the brains of the CX mice including ICAM-1, TNF-a and p-p65. Establishing the validity of the neuroinflammation phenotype in CS and the potential link with NE defects observed in CS-A cells using iPSC-derived or transdifferentiated neuronal CSA patient cell lines would be an interesting area of investigation.

Overall, our study has identified a new, non-canonical function of CSA in regulating NE integrity independent of its established role in the TC-NER pathway. This further reinforce the role played by NE dysfunction in various age-related conditions, outside of the known laminopathies. In CS-A cells, NE fragility and rupturing may contribute to the accumulation of DNA damage over time and to the activation of innate immune pathway activation, potentially contributing to inflammation in tissues such as the brain. Further understanding of the mechanisms behind the NE defects in CSA patient derived cells may help explaining some of the clinical phenotypes observed in CS patients and open new therapeutic avenues.

## Acknowledgments

We thank Dr. Sebastian Iben for providing the CS-A, WT(HA-CSA), CSB, and WT(HA-CSB) cell lines and the Flag-CSA-containing plasmid. We thank Dr. Kyle Roux for providing the LEMD2-GFP-containing plasmid. The authors gratefully acknowledge the Cambridge Advanced Imaging Center for their support and assistance with the use of Stellaris 5 confocal microscope in this work. This research was supported by the CIMR Microscopy Core Facility. In particular, we wish to thank Matthew Gratian and Mark Bowen for their advice and support in live imaging with the Incucyte. We would like to acknowledge Dr. Nicola Lawrence at the Gurdon Institute Imaging Facility for microscopy and image analysis support. The authors declare that they have no conflict of interest.

## Funding

D.Y. is currently funded by a Taiwan Cambridge Scholarship. A.L. and D.L were funded by a Sir Henry Dale Fellowship jointly funded by the Wellcome Trust and the Royal Society (Grant Number 206242/Z/17/Z. A.F.J.J. was supported by a FEBS Long-Term Fellowship as well as a Leverhulme Trust Early Career Fellowship and the Isaac Newton Trust.

## Supplementary Figures

**Figure S1:**
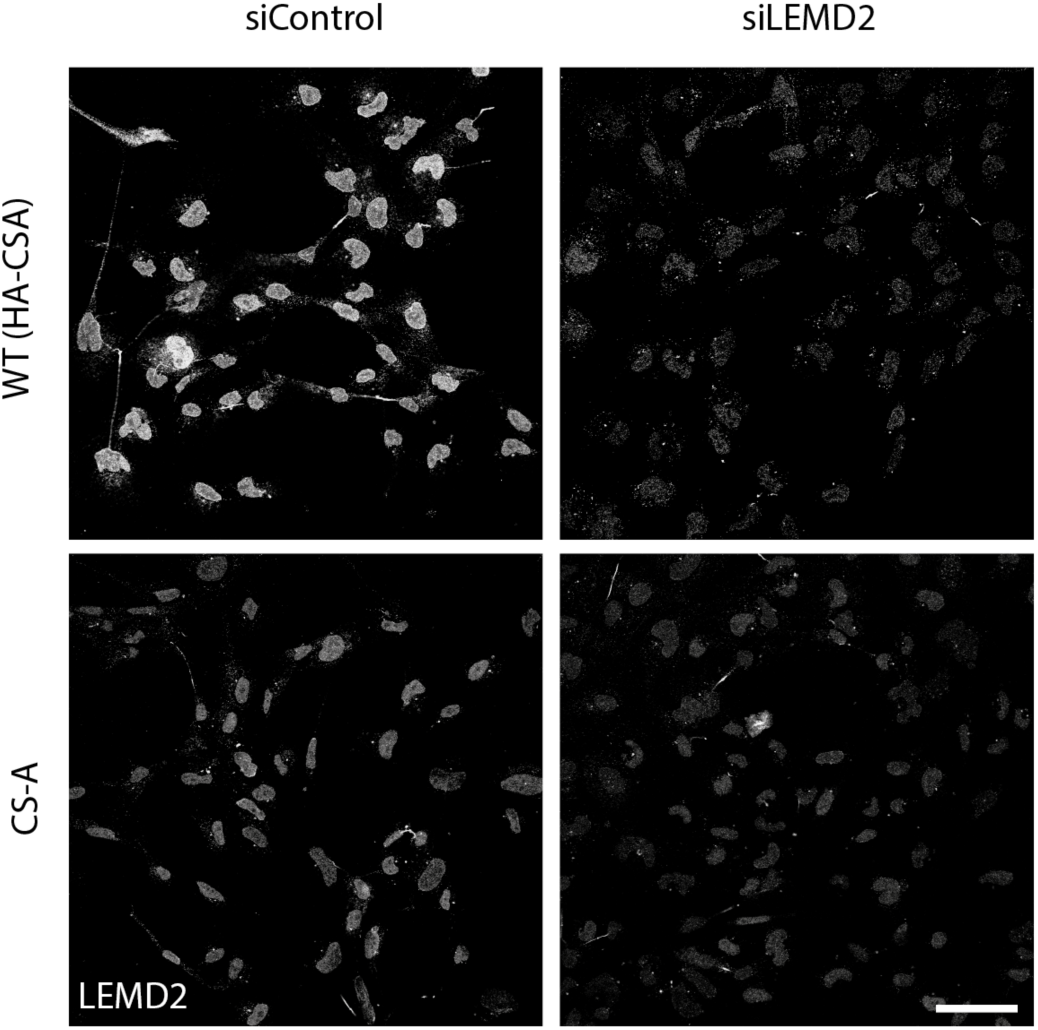
Assessing the specificity of the LEMD2 antibody. Immunofluorescence staining of LEMD2 in pre-extracted WT(HA-CSA) and CS-A cells following siRNA mediated knock down of LEMD2 (siLEMD2); scale bar: 50 µm.

## References

1. Rieckher M, Garinis GA, Schumacher B. Molecular pathology of rare progeroid diseases. Trends Mol Med. 2021 Sep;27(9):907–22.

2. Schnabel F, Kornak U, Wollnik B. Premature aging disorders: A clinical and genetic compendium. Clin Genet. 2021 Jan 29;99(1):3–28.

3. Carrero D, Soria-Valles C, López-Otín C. Hallmarks of progeroid syndromes: lessons from mice and reprogrammed cells. Dis Model Mech. 2016 Jul 1;9(7):719–35.

4. Tan WH, Baris H, Robson CD, Kimonis VE. Cockayne syndrome: The developing phenotype. Am J Med Genet A. 2005 Jun 1;135A(2):214–6.

5. Nardo T, Oneda R, Spivak G, Vaz B, Mortier L, Thomas P, et al. A UV-sensitive syndrome patient with a specific *CSA* mutation reveals separable roles for CSA in response to UV and oxidative DNA damage. Proceedings of the National Academy of Sciences. 2009 Apr 14;106(15):6209–14.

6. Troelstra C, van Gool A, de Wit J, Vermeulen W, Bootsma D, Hoeijmakers JHJ. ERCC6, a member of a subfamily of putative helicases, is involved in Cockayne’s syndrome and preferential repair of active genes. Cell. 1992 Dec;71(6):939–53.

7. Kleijer WJ, Laugel V, Berneburg M, Nardo T, Fawcett H, Gratchev A, et al. Incidence of DNA repair deficiency disorders in western Europe: Xeroderma pigmentosum, Cockayne syndrome and trichothiodystrophy. DNA Repair (Amst). 2008 May;7(5):744–50.

8. Marteijn JA, Lans H, Vermeulen W, Hoeijmakers JHJ. Understanding nucleotide excision repair and its roles in cancer and ageing. Nat Rev Mol Cell Biol. 2014 Jul 23;15(7):465–81.

9. Laugel V. Cockayne syndrome: The expanding clinical and mutational spectrum. Mech Ageing Dev. 2013 May;134(5–6):161–70.

10. Karikkineth AC, Scheibye-Knudsen M, Fivenson E, Croteau DL, Bohr VA. Cockayne syndrome: Clinical features, model systems and pathways. Ageing Res Rev. 2017 Jan;33:3–17.

11. Fousteri M, Mullenders LH. Transcription-coupled nucleotide excision repair in mammalian cells: molecular mechanisms and biological effects. Cell Res. 2008 Jan 1;18(1):73–84.

12. Saijo M. The role of Cockayne syndrome group A (CSA) protein in transcription-coupled nucleotide excision repair. Mech Ageing Dev. 2013 May;134(5–6):196–201.

13. Xu C, Min J. Structure and function of WD40 domain proteins. Protein Cell. 2011 Mar 6;2(3):202–14.

14. Sugita K, Suzuki N, Kojima T, Tanabe Y, Nakajima H, Hayashi A, et al. Cockayne Syndrome with Delayed Recovery of RNA Synthesis after Ultraviolet Irradiation but Normal Ultraviolet Survival. Pediatr Res. 1987 Jan;21(1):34–7.

15. Vessoni AT, Herai RH, Karpiak J V., Leal AMS, Trujillo CA, Quinet A, et al. Cockayne syndrome-derived neurons display reduced synapse density and altered neural network synchrony. Hum Mol Genet. 2016 Apr 1;25(7):1271–80.

16. Itoh T, Fujiwara Y, Ono T, Yamaizumi M. UVs syndrome, a new general category of photosensitive disorder with defective DNA repair, is distinct from xeroderma pigmentosum variant and rodent complementation group I. Am J Hum Genet. 1995 Jun;56(6):1267–76.

17. Scheibye-Knudsen M, Ramamoorthy M, Sykora P, Maynard S, Lin PC, Minor RK, et al. Cockayne syndrome group B protein prevents the accumulation of damaged mitochondria by promoting mitochondrial autophagy. Journal of Experimental Medicine. 2012 Apr 9;209(4):855–69.

18. de Waard H, de Wit J, Andressoo JO, van Oostrom CTM, Riis B, Weimann A, et al. Different Effects of *CSA* and *CSB* Deficiency on Sensitivity to Oxidative DNA Damage. Mol Cell Biol. 2004 Sep 1;24(18):7941–8.

19. Dechat T, Adam SA, Goldman RD. Nuclear lamins and chromatin: When structure meets function. Adv Enzyme Regul. 2009 Jan;49(1):157–66.

20. Taddei A, Hediger F, Neumann FR, Gasser SM. The Function of Nuclear Architecture: A Genetic Approach. Annu Rev Genet. 2004 Dec 1;38(1):305–45.

21. Crisp M, Liu Q, Roux K, Rattner JB, Shanahan C, Burke B, et al. Coupling of the nucleus and cytoplasm: Role of the LINC complex. J Cell Biol. 2006 Jan 2;172(1):41–53.

22. Bridger JM, Foeger N, Kill IR, Herrmann H. The nuclear lamina. FEBS Journal. 2007 Mar;274(6):1354–61.

23. Brachner A, Reipert S, Foisner R, Gotzmann J. LEM2 is a novel MAN1-related inner nuclear membrane protein associated with A-type lamins. J Cell Sci. 2005 Dec 15;118(24):5797–810.

24. Maurer M, Lammerding J. The Driving Force: Nuclear Mechanotransduction in Cellular Function, Fate, and Disease. Annu Rev Biomed Eng. 2019 Jun 4;21(1):443–68.

25. Isermann P, Lammerding J. Nuclear Mechanics and Mechanotransduction in Health and Disease. Current Biology. 2013 Dec;23(24):R1113–21.

26. Cohen TV, Hernandez L, Stewart CL. Functions of the nuclear envelope and lamina in development and disease. Biochem Soc Trans. 2008 Dec 1;36(6):1329–34.

27. Kalukula Y, Stephens AD, Lammerding J, Gabriele S. Mechanics and functional consequences of nuclear deformations. Nat Rev Mol Cell Biol. 2022 Sep 5;23(9):583–602.

28. Raab M, Gentili M, de Belly H, Thiam HR, Vargas P, Jimenez AJ, et al. ESCRT III repairs nuclear envelope ruptures during cell migration to limit DNA damage and cell death. Science (1979). 2016 Apr 15;352(6283):359–62.

29. Bell ES, Shah P, Zuela-Sopilniak N, Kim D, Varlet AA, Morival JLP, et al. Low lamin A levels enhance confined cell migration and metastatic capacity in breast cancer. Oncogene. 2022 Sep 2;41(36):4211–30.

30. Srivastava N, Nader GP de F, Williart A, Rollin R, Cuvelier D, Lomakin A, et al. Nuclear fragility, blaming the blebs. Curr Opin Cell Biol. 2021 Jun;70:100–8.

31. Young AM, Gunn AL, Hatch EM. BAF facilitates interphase nuclear membrane repair through recruitment of nuclear transmembrane proteins. Mol Biol Cell. 2020 Jul 15;31(15):1551–60.

32. Maciejowski J, Hatch EM. Nuclear Membrane Rupture and Its Consequences. Annu Rev Cell Dev Biol. 2020 Oct 6;36(1):85–114.

33. Halfmann CT, Sears RM, Katiyar A, Busselman BW, Aman LK, Zhang Q, et al. Repair of nuclear ruptures requires barrier-to-autointegration factor. Journal of Cell Biology. 2019 Jul 1;218(7):2136–49.

34. von Appen A, LaJoie D, Johnson IE, Trnka MJ, Pick SM, Burlingame AL, et al. LEM2 phase separation promotes ESCRT-mediated nuclear envelope reformation. Nature. 2020 Jun 4;582(7810):115–8.

35. Gu M, LaJoie D, Chen OS, von Appen A, Ladinsky MS, Redd MJ, et al. LEM2 recruits CHMP7 for ESCRT-mediated nuclear envelope closure in fission yeast and human cells. Proceedings of the National Academy of Sciences. 2017 Mar 14;114(11).

36. Janssen AFJ, Breusegem SY, Larrieu D. Current Methods and Pipelines for Image-Based Quantitation of Nuclear Shape and Nuclear Envelope Abnormalities. Cells. 2022 Jan 20;11(3):347.

37. Epanchintsev A, Costanzo F, Rauschendorf MA, Caputo M, Ye T, Donnio LM, et al. Cockayne’s Syndrome A and B Proteins Regulate Transcription Arrest after Genotoxic Stress by Promoting ATF3 Degradation. Mol Cell. 2017 Dec;68(6):1054–1066.e6.

38. Martins F, Sousa J, Pereira CD, da Cruz e Silva OAB, Rebelo S. Nuclear envelope dysfunction and its contribution to the aging process. Aging Cell. 2020 May 15;19(5).

39. Gentili M, Lahaye X, Nadalin F, Nader GPF, Puig Lombardi E, Herve S, et al. The N-Terminal Domain of cGAS Determines Preferential Association with Centromeric DNA and Innate Immune Activation in the Nucleus. Cell Rep. 2019 Feb;26(9):2377–2393.e13.

40. Ulbert S, Antonin W, Platani M, Mattaj IW. The inner nuclear membrane protein Lem2 is critical for normal nuclear envelope morphology. FEBS Lett. 2006 Nov 27;580(27):6435– 41.

41. Moser B, Basílio J, Gotzmann J, Brachner A, Foisner R. Comparative Interactome Analysis of Emerin, MAN1 and LEM2 Reveals a Unique Role for LEM2 in Nucleotide Excision Repair. Cells. 2020 Feb 18;9(2):463.

42. Alupei MC, Maity P, Esser PR, Krikki I, Tuorto F, Parlato R, et al. Loss of Proteostasis Is a Pathomechanism in Cockayne Syndrome. Cell Rep. 2018 May;23(6):1612–9.

43. Qiang M, Khalid F, Phan T, Ludwig C, Scharffetter-Kochanek K, Iben S. Cockayne Syndrome-Associated CSA and CSB Mutations Impair Ribosome Biogenesis, Ribosomal Protein Stability, and Global Protein Folding. Cells. 2021 Jun 28;10(7):1616.

44. Maninova M, Caslavsky J, Vomastek T. The assembly and function of perinuclear actin cap in migrating cells. Protoplasma. 2017 May 18;254(3):1207–18.

45. Tojkander S, Gateva G, Lappalainen P. Actin stress fibers – assembly, dynamics and biological roles. J Cell Sci. 2012 Jan 1;

46. Hatch EM, Hetzer MW. Nuclear envelope rupture is induced by actin-based nucleus confinement. Journal of Cell Biology. 2016 Oct 10;215(1):27–36.

47. Janssen A, Marcelot A, Breusegem S, Legrand P, Zinn-Justin S, Larrieu D. The BAF A12T mutation disrupts lamin A/C interaction, impairing robust repair of nuclear envelope ruptures in Nestor–Guillermo progeria syndrome cells. Nucleic Acids Res. 2022 Sep 9;50(16):9260–78.

48. Decout A, Katz JD, Venkatraman S, Ablasser A. The cGAS–STING pathway as a therapeutic target in inflammatory diseases. Nat Rev Immunol. 2021 Sep 8;21(9):548–69.

49. López-Otín C, Blasco MA, Partridge L, Serrano M, Kroemer G. Hallmarks of aging: An expanding universe. Cell. 2023 Jan;186(2):243–78.

50. De Vos WH, Houben F, Kamps M, Malhas A, Verheyen F, Cox J, et al. Repetitive disruptions of the nuclear envelope invoke temporary loss of cellular compartmentalization in laminopathies. Hum Mol Genet. 2011 Nov 1;20(21):4175–86.

51. Prissette M, Fury W, Koss M, Racioppi C, Fedorova D, Dragileva E, et al. Disruption of nuclear envelope integrity as a possible initiating event in tauopathies. Cell Rep. 2022 Aug;40(8):111249.

52. Paonessa F, Evans LD, Solanki R, Larrieu D, Wray S, Hardy J, et al. Microtubules Deform the Nuclear Membrane and Disrupt Nucleocytoplasmic Transport in Tau-Mediated Frontotemporal Dementia. Cell Rep. 2019 Jan;26(3):582–593.e5.

53. Sohn C, Ma J, Ray WJ, Frost B. Pathogenic tau decreases nuclear tension in cultured neurons. Frontiers in Aging. 2023 Jan 23;4.

54. Caravia XM, Ramirez-Martinez A, Gan P, Wang F, McAnally JR, Xu L, et al. Loss of function of the nuclear envelope protein LEMD2 causes DNA damage–dependent cardiomyopathy. Journal of Clinical Investigation. 2022 Nov 15;132(22).

55. Marbach F, Rustad CF, Riess A, Đukić D, Hsieh TC, Jobani I, et al. The Discovery of a LEMD2-Associated Nuclear Envelopathy with Early Progeroid Appearance Suggests Advanced Applications for AI-Driven Facial Phenotyping. The American Journal of Human Genetics. 2019 Apr;104(4):749–57.

56. Abdelfatah N, Chen R, Duff HJ, Seifer CM, Buffo I, Huculak C, et al. Characterization of a Unique Form of Arrhythmic Cardiomyopathy Caused by Recessive Mutation in LEMD2. JACC Basic Transl Sci. 2019 Apr;4(2):204–21.

57. Chen R, Buchmann S, Kroth A, Arias-Loza AP, Kohlhaas M, Wagner N, et al. Mechanistic Insights of the LEMD2 p.L13R Mutation and Its Role in Cardiomyopathy. Circ Res. 2023 Jan 20;132(2).

58. Ross JA, Arcos-Villacis N, Battey E, Boogerd C, Orellana CA, Marhuenda E, et al. Lem2 is essential for cardiac development by maintaining nuclear integrity. Cardiovasc Res. 2023 Apr 17;

59. Chen CY, Chi YH, Mutalif RA, Starost MF, Myers TG, Anderson SA, et al. Accumulation of the Inner Nuclear Envelope Protein Sun1 Is Pathogenic in Progeric and Dystrophic Laminopathies. Cell. 2012 Apr;149(3):565–77.

60. Shah PP, Santini GT, Shen KM, Jain R. InterLINCing Chromatin Organization and Mechanobiology in Laminopathies. Curr Cardiol Rep. 2023 May 13;25(5):307–14.

61. Chai RJ, Werner H, Li PY, Lee YL, Nyein KT, Solovei I, et al. Disrupting the LINC complex by AAV mediated gene transduction prevents progression of Lamin induced cardiomyopathy. Nat Commun. 2021 Aug 5;12(1):4722.

62. Yum S, Li M, Fang Y, Chen ZJ. TBK1 recruitment to STING activates both IRF3 and NF-κB that mediate immune defense against tumors and viral infections. Proceedings of the National Academy of Sciences. 2021 Apr 6;118(14).

63. Andrade B, Jara-Gutiérrez C, Paz-Araos M, Vázquez MC, Díaz P, Murgas P. The Relationship between Reactive Oxygen Species and the cGAS/STING Signaling Pathway in the Inflammaging Process. Int J Mol Sci. 2022 Dec 2;23(23):15182.

64. Kellogg DL, Kellogg DL, Musi N, Nambiar AM. Cellular Senescence in Idiopathic Pulmonary Fibrosis. Curr Mol Biol Rep. 2021 Sep 12;7(3):31–40.

65. Li X, Li C, Zhang W, Wang Y, Qian P, Huang H. Inflammation and aging: signaling pathways and intervention therapies. Signal Transduct Target Ther. 2023 Jun 8;8(1):239.

66. Kajitani GS, Brace L, Trevino-Villarreal JH, Trocha K, MacArthur MR, Vose S, et al. Neurovascular dysfunction and neuroinflammation in a Cockayne syndrome mouse model. Aging. 2021 Oct 15;13(19):22710–31.

